# Spatial, temporal and sex specific mitochondrial dynamic changes in severe controlled cortical impact mouse model of traumatic brain injury

**DOI:** 10.64898/2026.04.20.719702

**Authors:** Hemendra J. Vekaria, Chirayu D. Pandya, Paresh Prajapati, Elika Z. Moallem, Gopal V. Velmurugan, W. Brad Hubbard, Adam D. Bachstetter, Patrick G. Sullivan

## Abstract

Traumatic brain injury (TBI) triggers complex and evolving secondary cascades that disrupt mitochondrial homeostasis and contribute to progressive neurodegeneration. Although mitochondrial impairment is a well-recognized driver of post-traumatic pathology, the spatial and temporal progression of mitochondrial dysfunction, particularly in regions distal to the injury site, remains poorly defined, and potential sex-specific responses remain understudied. Here, we performed a comprehensive mitochondrial-focused analysis in a mouse model of controlled cortical impact (CCI), quantifying mtDNA copy number (mtDNA-CN), mitochondrial gene expression, and protein markers regulating biogenesis, transcription, electron transport chain integrity, and mitophagy. Mitochondrial profiles were assessed across four brain regions (cortex at 2, 4, and 6 mm from the injury epicenter, and hippocampus) at four time points (6h, 12h, 24h, and 48h) in both female and male C57BL/6J mice. While mtDNA content exhibited only modest and region-restricted reduction, particularly near the injury core, transcriptional and protein-level changes were far more pronounced and sex-divergent. Females displayed extensive early cortical gene activation followed by widespread hippocampal suppression at 48 h across mitochondrial dynamics, OXPHOS, transcriptional regulation, and biogenesis pathways, accompanied by 48h in PGC-1α, TFAM, and NDUFS1. In contrast, males showed minimal transcriptional disruption but demonstrated delayed compensatory increases in TFAM, NDUFS1, and p62 protein levels, suggesting activation of mitochondrial maintenance and recovery programs. These spatially and temporally distinct responses reveal fundamental sex-specific vulnerabilities in mitochondrial regulation after TBI. Together, our findings provide a direction to an integrated mitochondrial landscape of early post-injury events and identifies critical windows and pathways that may support sex-specific therapeutic targeting to restore mitochondrial function after TBI.

## Introduction

Traumatic brain injury (TBI) leads to a complex cascade of secondary pathology that propagates temporally and spatially, affecting regions which are directly impacted and also distal to the primary injury site ^1,2^. Among these secondary processes, mitochondrial dysfunction is a proven central driver of neuronal vulnerability, with bioenergetic failure contributing to synaptic impairment and progressive neuronal loss ^3^, changes that are particularly well characterized in controlled cortical impact (CCI) models. Within hours of injury, cortical mitochondria exhibit acute impairments in oxidative phosphorylation, loss of mitochondrial membrane potential and excessive Ca^2+^ uptake driving opening of the mitochondrial permeability transition pore (mPTP) and rapid ATP depletion ^4-6^. Elevated reactive oxygen species (ROS) production further oxidizes mitochondrial proteins and lipids, amplifying bioenergetic collapse ^7,8^. Over subsequent days, mitochondrial respiratory chain complexes, particularly complexes I and IV remain depressed, while dysregulated fission/fusion dynamics, including DRP1 hyperactivation and OPA1 loss, impair structural integrity and cristae function ^9-11^. These deficits progress heterogeneously across brain regions, with the hippocampus displaying delayed reductions in spare respiratory capacity and synaptic mitochondrial function while the cerebellum shows comparatively resilient metabolic trajectories ^12^. Subacute (0h - 24h) and acute (24h – 1 week) phases of CCI are marked by persistent inflammation-driven mitochondrial fragmentation, impaired mitochondrial biogenesis linked to PGC-1α suppression ^13,14^.

Time-dependent alterations in mitochondrial gene regulation further shape the trajectory of neuroenergetic failure and recovery. In the acute and sub-acute phase, mitochondrial respiratory chain genes such as ND1, ND5/ND6, Ndufs1, Cox I, and ATP6 are rapidly downregulated, reflecting impaired electron transport and ATP synthesis ^15,16^. Concurrently, increased UCP2 expression represents a compensatory attempt to limit oxidative stress by mild uncoupling ^17^. Mitophagy regulators Pink1 and PARK2 are upregulated early to clear damaged mitochondria ^18,19^, while fission–fusion balance shifts toward fragmentation through elevated Fis1 and DNM1 and reduced Mfn1/Mfn2 ^20^. Over days to weeks, metabolic recovery depends on mitochondrial biogenesis pathways: Ppargc1α, Sirt1, Tfam, Nrf1, Gtf2h1, and Nfe2l2 gradually increase to restore mitochondrial mass and antioxidant responses ^21,22^. Chronic stages often show persistent deficits in energy sensing, marked by dysregulation of Prkaa2 (AMPKα2) and reduced NOS3, contributing to long-term metabolic insufficiency and neurodegeneration ^23^.

Despite extensive characterization of mitochondrial dysfunction following CCI, the spatial boundaries of these deficits remain poorly defined. Most studies rely on whole-brain homogenates or broad cortical samples, obscuring how mitochondrial pathology extends from the injury core into surrounding tissue. However, without systematic assessment of how far mitochondrial dysfunction propagates from the primary lesion, it remains unclear which regions would benefit from mitochondria-directed intervention.

To address this gap, we systematically evaluated mitochondrial gene expression, protein abundance, and mtDNA copy number (mtDNA-CN) at the injury core and at defined intervals into surrounding tissue across acute and subacute timepoints. We further examined whether males and females exhibit distinct mitochondrial responses. Together, these studies define the spatiotemporal boundaries of mitochondrial dysfunction after CCI and identify potential targets for region-specific intervention.

## Materials and Methods

### Animals

Wild-type, 2-3 months old C57BL/6J male and female mice (Jackson Laboratorie) were used for the study. The average body weights were around 25 g (male) or 20 g (female). Animal protocols were approved by University of Kentucky, Institutional Animal Care and Use Committee (IACUC) and also by the United States Veterans Affairs Animal Component of Research Protocol. All animal procedures complied with ARRIVE (Animal Research: Reporting of In Vivo Experiments) guidelines for *Laboratory Animals*. Animals were housed at 5 per cage maximum and under a 14h and 10h, light and dark cycle respectively. The mice had constant access to ad libitum balanced diet and water. Upon arrival, the animals were acclimatized for at least one week before performing any surgeries. Sham or surgery procedures on animals were carried out in a randomized manner with equal distribution of groups in each experimental set.

### Controlled cortical impact surgery in mice

The mice were operated on for sham (craniotomy only) or controlled cortical impact (CCI) procedures as done in previous studies ^24^. The mice heads were shaved under 4% isoflurane anesthesia and were ear-barred onto a stereotaxic frame. Isoflurane inhalation (2.5%) via nose cone and warm body temperature using a heating pad was constantly maintained throughout the procedure. 50µl Bupivacaine (#77614, Covetrus) was injected subcutaneously at the site of injury. The skin at the site of injury was disinfected by wiping with iodine and alcohol. Approximately 1cm vertical cut was made over the head starting between the eyes and extending towards posterior to expose the bragma and lambda points on the skull. Using a motorized Dremel, a circular craniotomy approximately 5 mm in diameter was carefully performed on left side to the sagittal suture, positioned between bregma and lambda. The trimmed skull was carefully removed without touching the dura. The animal with the stereotaxis apparatus were mounted on the impactor (Precision Systems and Instrumentation). For the CCI group, a 3-mm diameter tip was used to deliver a cortical impact at a velocity of 3.5 m/s, a dwell time of 500 ms, and an impact depth of 1.0 mm. As per previous reports, these conditions generate medium to severe types of brain injury ^24^. After the impact, hemostatic dressing was placed over the exposed cortex covering the injury site. Immediately the skin was pooled with forceps, and the injury site was closed with staples. The mice were placed under observation in a clean cage kept over a heating pad maintained at 37 °C until consciousness was regained. No mice showed pain or distress symptoms after surgery. Mice were euthanized at respective times, 6h, 12h, 24h, and 48h after surgery using the CO2 chamber and immediately decapitated for brain tissue collection.

### Brain tissue collection

The study was performed by dividing all the mice into 8 sets, separated based on sex and experimental timepoint, 4 sets each of males and females (Fig.1A). Each of the 4 sets were separated based on the experimental timepoint of 6h, 12h, 24h, and 48h after CCI. At each termination time point, the mice were sacrificed, heads were decapitated; brains were removed quickly and placed on an ice cooled metal block. As shown in figure 1A, the cortex and hippocampus tissues from the ipsilateral side of the injury were removed. Each cortex was further separated into 3 regions based on its distance from the site of injury, by using 3 different punches of 2mm, 4mm, and 6mm diameters. The immediate cortex under the impact was extracted using a 2mm punch. The immediate penumbra cortex was extracted using a 4mm punch followed by the distal cortex penumbra using a 6mm punch. Each separated cortical and hippocampal tissue was transferred to a 1.5ml centrifuge tube containing ice cold 250µl phosphate buffered saline. The tissues were homogenized quickly using a handheld motorized micro pestle (Fisher scientific #12-141-361). Each homogenate was distributed into three 1.5ml tubes. 100µl was transferred to a tube containing 750µl Trizol (ThermoFisher #10296028) for RNA isolation, 100µl was transferred to a 1.5ml tube containing a mix of 10x RIPA and 1µl of 100X protease inhibitor cocktail (Sigma #P8340) which were used for western blot analysis (Fig.1B). The remaining 50µl was used for DNA isolation. The samples collected were frozen immediately on dry ice and were stored at –80C till the assays were performed.

**FIG. 1.**
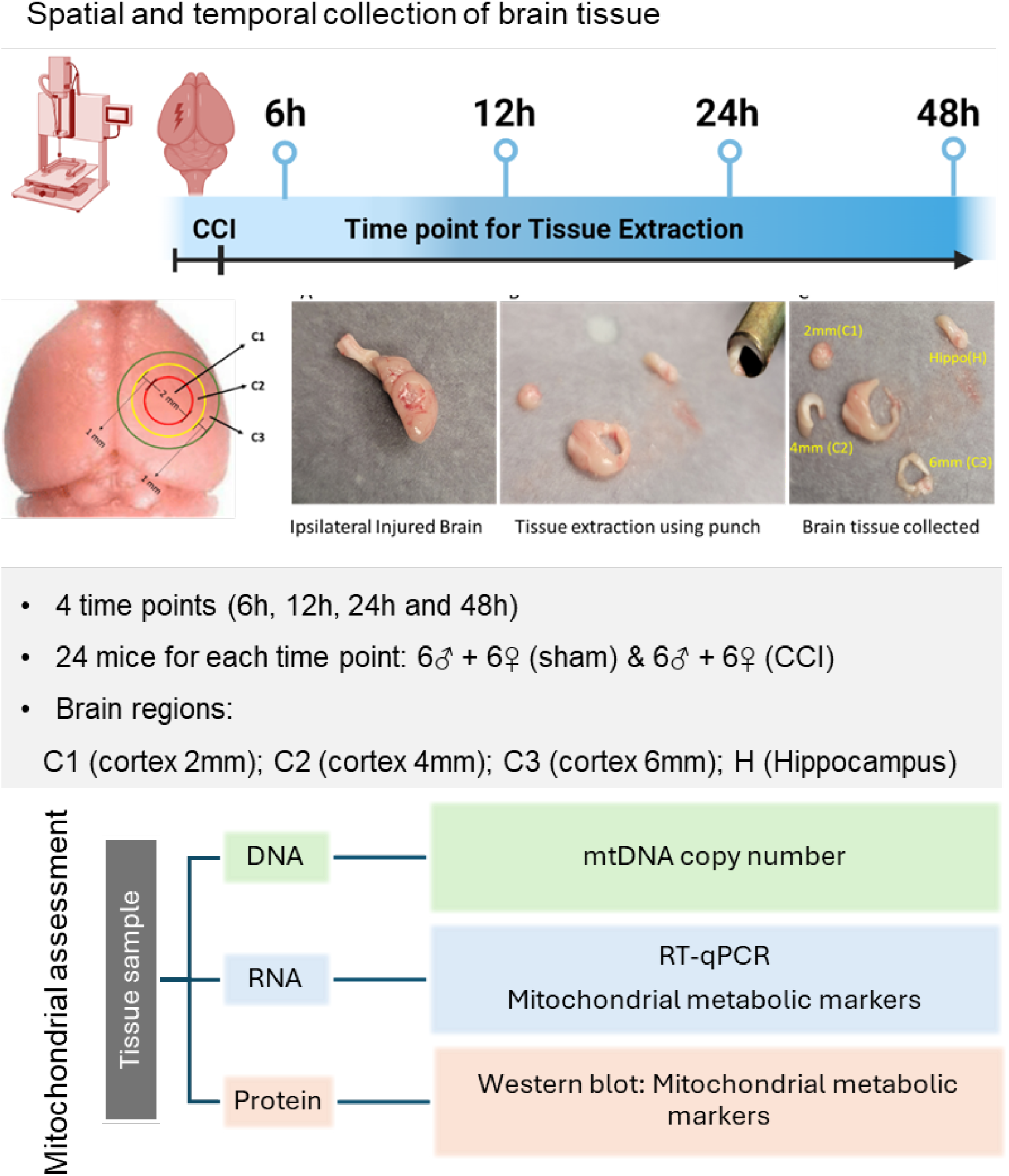
Experimental design and spatial–temporal tissue sampling following controlled cortical impact (CCI). Schematic overview of the study workflow. Adult male and female mice underwent controlled cortical impact (CCI) or sham surgery. Brains were collected at 6, 12, 24, and 48 h post-injury. For molecular analyses, tissue punches were obtained from cortex at 2 mm, 4 mm, and 6 mm from the injury epicenter, as well as from the ipsilateral hippocampus. Mitochondrial DNA (mtDNA) copy number, mitochondrial gene expression, and protein markers of mitochondrial biogenesis and quality control (PGC-1α, Tfam, Ndufs1, p62) were quantified in each region across all timepoints. This design enabled spatially and temporally resolved characterization of post-TBI mitochondrial responses in both sexes.

### Mitochondrial DNA copy number (mtDNA-CN) estimation

The homogenate fractions collected for DNA isolation were thawed on ice. DNA was purified with DNeasy Blood and Tissue Kit (Qiagen #69504) as per the standard protocol and DNA was quantified using NonoDrop-8000 (Thermo Scientific). Quantitative PCR was done using PCR PowerTrack SYBR (Thermo Fisher Scientific #A46109) assay on Quantstudio-7 (Applied Biosystems). 5 ng DNA was used as a template in a 10µl system for quantification of relative mtDNA content. qPCR analysis of a gene fragment of ND1, a mitochondrial gene, was carried out using primers, ND1-forward (5′-TGAATCCGAGCATCCTACC-3′) and ND1-reverse (5′-ATTCCTGCTAGGAAAATTGG-3′). The mitochondrial DNA amplification was normalized to that of a section of a nuclear-encoded β-actin gene. The primers used were Actin-forward (5′-GGGATGTTTGCTCCAACCAA-3′) and Actin-reverse (5′-GCGCTTTTGACTCAGGATTTAA-3′) The ΔCt values obtained from qPCR were used to calculate differences between sham vs CCI groups and the data were analyzed using Graph-Pad prism as mentioned in statistical analysis.

### Quantitative Reverse Transcriptase Polymerase Chain Reaction (qRT-PCR)

Trizol containing RNA sample tubes were thawed on ice and were topped with 150µl PBS to make up total volume to 1ml. RNA was isolated using the standard trizol method. The purified RNA was quantified using NonoDrop-8000 (ThermoFisher Scientific). 1µg of RNA was then reverse transcribed with a qScript Ultra SuperMix (Quanta Biosciences, #95217) according to the manufacturer’s recommendations. The cDNA was amplified and analyzed using Taqman Gene Expression assay probes (FAM or VIC) (**Table 1**) (ThermoFisher Scientific, # 4453320) in combination with TaqMan™ Fast Advanced Master Mix (ThermoFisher Scientific, # 4444963). The amplification cycles were run on a 384 well QuantStudio 7 real time PCR machine (Thermo Fisher Scientific, Waltham, MA). The Actb and Rn18s housekeeping genes were used as endogenous controls. Relative expression was estimated using the 2^ΔΔ-Ct^ method. The relative expression of each gene was normalized to the respective sham control for each respective injury. The data are reported as fold change. For the panel of mitochondria-dynamics-related genes, expression values were additionally converted to z-scores relative to sham controls. Graphs were generated using R. Taqman primers were purchased through ThermoFisher Scientific and are listed in Table 1.

**Table 1.** Primer sequences for mitochondrial DNA and gene expression analyses. List of primers used for quantitative PCR analyses. The table includes forward and reverse sequences for mtDNA targets (e.g., mt-ND1), nuclear reference genes, and the 20 mitochondrial-related genes categorized under mitochondrial dynamics, mitochondrial function/ETC components, and transcriptional regulators. All primers were validated for specificity and efficiency (90–110%) prior to use.

| S. No. | Gene Symbol | Gene Name | Assay ID |
| --- | --- | --- | --- |
| 1 | Pink1 | PTEN-induced putative kinase 1 | Mm00550827 |
| 3 | Mfn2 | Mitofusin 2 | Mm00500120 |
| 4 | Mfn1 | Mitofusin 1 | Mm00612599_m1 |
| 5 | Gtf2h1 | General transcription factor IIH subunit 1 | Mm00500417_m1 |
| 6 | Fis1 | Fission protein 1 | Mm00481580 |
| 7 | DNM1 | Dynamin-related GTPase Dnm1 | Mm01342903_m1 |
| 8 | Ucp2 | Uncoupling protein 2 | Mm00627599_m1 |
| 9 | Ndufs1 | NADH:Ubiquinone Oxidoreductase Core Subunit S1 | Mm005236040_m1 |
| 10 | ND6/ND5 | NADH-ubiquinone oxidoreductase chain 6 protein/5 | Mm04225325_g1 |
| 11 | ND1 | NADH-ubiquinone oxidoreductase chain 1 | Mm04225274_s1 |
| 12 | COX1 | Cytochrome c oxidase I | Mm04225243_g1 |
| 13 | ATP6 | ATP Synthase Membrane Subunit 6 | Mm03649417_g1 |
| 14 | TFAM | Transcription Factor A, Mitochondrial | Mm00447485_m1 |
| 14 | Dnm1 | Dynamin 1 | Mm01342903_m1 |
| 15 | Sirt1 | Sirtuin 1 | Mm01168521_m1 |
| 16 | Prkaa2 | Protein Kinase AMP-Activated Catalytic Subunit Alpha 2 | Mm01264789_m1 |
| 17 | Ppargc1 | Peroxisome proliferator-activated receptor-gamma coactivator 1-alpha (PGC-1α) | Mm01208835_m1 |
| 18 | Nrf1 | Nuclear respiratory factor 1 | Mm01135606_m1 |
| 19 | NOS3 | Nitric oxide synthase 3 | Mm00435217_m1 |
| 20 | Nfe2l2 (Nrf2) | Nuclear factor erythroid 2-related factor 2 | Mm00477784_m1 |
| 21 | 18sRNA | 18S ribosomal RNA | Mm03928990_g1 |

### OXPHOS Protein Expression

Changes in protein levels were determined using Western Blot analysis as per standard protocols. The proteins involved in mitochondrial biogenesis and oxidative phosphorylation (OXPHOS) were analyzed and quantified. The protein content of the reserved protein homogenates was determined using BCA protein estimation kit (ThermoFisher #23225). The protein samples were normalized with 1X RIPA and were mixed with XT sample buffer (Bio-Rad #1610791). The proteins were separated on 4–12% Bis-Tris criterion gel (Bio-Rad #3450125) at 150V. Western transfer was done using Bio-Rad Trans-blot Turbo Transfer system onto a nitrocellulose membrane. The primary antibodies used were PGC-1α, (1:1000, Proteintech #66369); TFAM, (1:1000, Abcam #ab131607); NDUFS1 (1:1000, Abcam #ab169540); p62 (1:1000, Abcam #ab91526) and β-actin (1:10,000, Proteintech #20536-1-AP). The fluorescent secondary antibodies used were goat anti-mouse (1:10,000, LI-COR #926-68070) or goat anti-rabbit (1:10,000, LI-COR #926-32211). The blots were scanned with Odyssey CLx Imaging System (Li-COR). Densitometric analysis was performed using ImageJ (NIH, Bethesda, MD, USA), with target protein expression normalized to β-actin.

### Statistical Analyses

All statistical analyses for mtDNA and protein expression were performed using GraphPad Prism (GraphPad Software Inc., La Jolla, CA, USA). For each time point and sex, relative expression values were converted to a percentage of their respective sham controls. Data for males and females were analyzed separately. To assess injury effects within each tissue region, data were analyzed using a two-way repeated measures analysis of variance (ANOVA), assuming sphericity. Tukey’s post hoc multiple comparisons test was applied for hypothesis testing.

Gene expression data were organized by brain region, time point after injury, sex, and injury status (CCI versus sham). For each gene at each region-time combination, we compared log2-transformed expression values between CCI and sham groups using Welch’s t-test, which does not assume equal variances between groups. The Benjamini-Hochberg false discovery rate (FDR) correction within each region-time stratum was applied to control for multiple comparisons. This approach adjusts for testing multiple genes while accounting for the nested experimental structure. Genes were considered significantly different between CCI and sham groups at FDR < 0.05. For visualization, we calculated a signed Z-score for each comparison to represent both the magnitude and direction of effect. The signed Z-score was computed as sign(log2FC) × Z, where Z is the normal quantile corresponding to the two-tailed p-value from the t-test and log2FC is the log2 fold change (CCI mean minus sham mean). This metric provides a standardized effect size that incorporates statistical significance and retains directionality, with positive values indicating upregulation in CCI relative to sham and negative values indicating downregulation. Results are presented as bubble plots with genes on the y-axis and time points on the x-axis, faceted by brain region (and sex were analyzed separately). Bubble color represents the signed Z-score (blue for downregulation, red for upregulation), bubble size reflects FDR significance thresholds (larger bubbles indicate stronger statistical significance), and black outlines denote comparisons reaching FDR < 0.05. Genes were grouped by functional category (mitochondrial dynamics, mitochondrial function, transcriptional regulation) based on known biological roles. All analyses were conducted in R version 4.5.0 using the tidyverse suite of packages. Plots were generated using ggplot2 with consistent Z-score scaling across all panels to enable direct comparisons.

## Results

### Mitochondrial DNA Copy Number (mtDNA-CN) Exhibits Limited Spatial and Temporal Changes After CCI

To assess mitochondrial content following traumatic brain injury, mtDNA-CN was quantified in cortex (2, 4, and 6 mm from the injury epicenter) and hippocampus at 6h, 12h, 24h, and 48h post-CCI in male and female mice (Fig.2). Comparing the sham vs CCI differences, Overall, mtDNA levels remained largely stable in the cortex penumbra and hippocampus following injury. Significant differences were observed in primary injured 2mm cortex. Though both male and female showed decreased trend in the mtDNA content from 6 to 24h, the percent changes were quite significant in male at 6h but female mice didn’t show a significant difference. In male mice, mtDNA-CN was significantly reduced in the 2-mm cortex at both 6 h (Fig.2A: F (3, 70) = 6.453; p = 0.044) and 24 h (Fig.2A: F (3, 70) = 6.453; p = 0.042) post-CCI, indicating an early and transient mitochondrial loss near the injury site. No significant mtDNA changes were detected in the 4-mm (Fig.2B) or 6-mm (Fig.2B) cortex or in the hippocampus across any time point (Fig.2D). In female mice, mtDNA-CN remained unchanged across all cortical regions and time points (Fig.2A-C), with hippocampal mtDNA levels (Fig.2D) also showing no differences relative to sham. mtDNA-CN findings indicate that CCI induces only modest and region-restricted reductions in mitochondrial DNA content, with males showing slightly more susceptibility near injury epicenter.

**FIG. 2.**
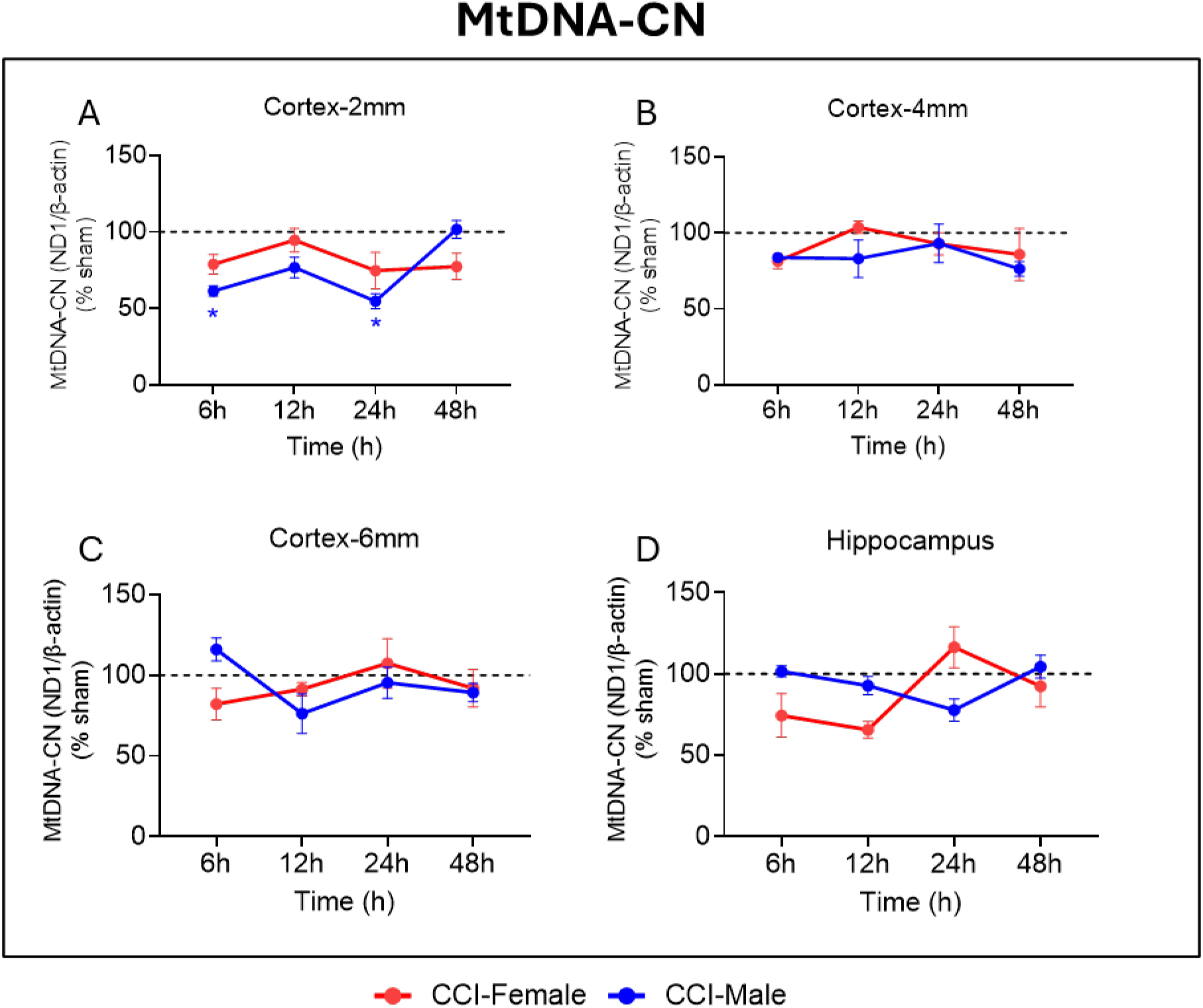
MtDNA-CN changes following CCI in female and male mice. MtDNA-CN were quantified in the **(A)** cortex-2 mm, **(B)** cortex-4 mm, **(C)** cortex-6 mm regions relative to the impact site, and **(D)** hippocampus at 6, 12, 24, and 48 h following CCI in female (red) and male (blue) mice. Data are expressed as relative expression (% of sham), with dotted horizontal lines indicating the sham reference level across all time points and regions. Data are presented as mean ± SEM, with individual biological replicates shown (n = 5–6 per group). Statistical significance is indicated as *p ≤ 0.05, and **p ≤ 0.01, compared with sham controls.

### Mitochondrial Gene Expression Following CCI Shows Pronounced Sex Differences

We examined expression of 20 genes involved in contrloling mitochondrial dynamics, mitochondrial function, and transcriptional regulation across four brain regions (cortex 2, 4, and 6 mm from the injury epicenter, and hippocampus) at four time points (6h, 12h, 24h, and 48h) following CCI (Fig.3). Of 640 total comparisons between CCI and sham groups, 27 reached statistical significance (FDR < 0.05). Strikingly, 26 of these significant changes (96%) occurred in female mice (Fig.3A), with only a single significant change observed in males (Fig.3B) (Ndufs1 downregulation in cortex-2mm at 12 h, FDR = 0.047). The most robust transcriptional response occurred in the female hippocampus at 48 h post-injury, where 16 of 20 genes showed significant alterations, all downregulated relative to sham. The mitochondrial dynamics genes Pink1 (FDR = 4.9×10^−4^), PARK2 (FDR = 9.4×10^−4^), Mfn2 (FDR = 5.8×10^−4^), Mfn1 (FDR = 7.6×10^−3^), Gff2h1 (FDR = 8.4×10^−3^), and Fis-I (FDR = 0.013) were all significantly reduced. Mitochondrial function genes likewise showed widespread downregulation, including Cox I (FDR = 1.0×10^−3^), ATP6 (FDR = 0.029), ND1 (FDR = 0.046), and Ndufs1 (FDR = 0.013). Transcriptional regulators Ppargc1 (FDR = 9.0×10^−4^), Prkaa2 (FDR = 1.0×10^−3^), Tfam (FDR = 1.6×10^−3^), Nrf1 (FDR = 4.2×10^−3^), and Sirt-I (FDR = 0.013) were similarly suppressed. In contrast to the late hippocampal suppression, the female cortex-4mm region exhibited significant upregulation of multiple mitochondrial genes at 12 h post-injury. Mitochondrial function genes Cox I (log2FC = 0.37, FDR = 0.021), ND1 (log2FC = 0.41, FDR = 0.027), and ATP6 (log2FC = 0.13, FDR = 0.034) showed increased expression. Mitochondrial dynamics genes Pink1 (FDR = 0.035), PARK2 (FDR = 0.035), Mfn1 (FDR = 0.027), DNM1 (FDR = 0.027), and Fis-I (FDR = 0.037) were also upregulated, along with the transcriptional regulator Prkaa2 (FDR = 0.027). Additionally, Nfe212 showed early upregulation in the hippocampus at 6 h (FDR = 0.038). In cortex-2mm, two genes showed significant changes in females: Ndufs1 was downregulated at both 12 h (FDR = 0.046) and 48 h (FDR = 3.3×10^−3^). In males, cortex-2mm showed the only significant change detected in male mice, with Ndufs1 downregulation at 12 h (FDR = 0.047). Neither female nor male mice exhibited significant gene expression changes in the cortex-6mm region at any time point examined, suggesting that transcriptional alterations were localized to regions closer to the injury epicenter and the hippocampus.

**FIG. 3.**
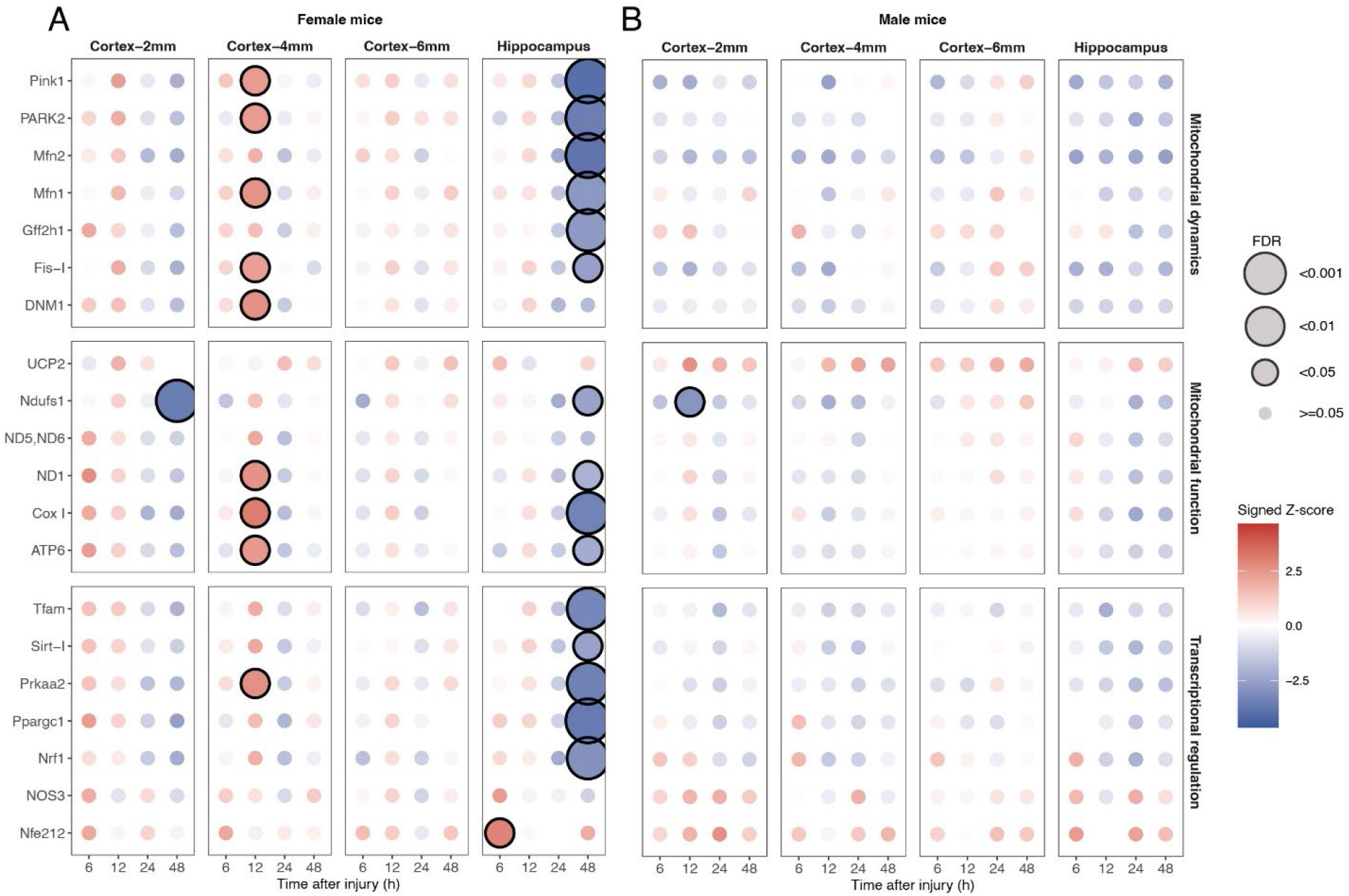
Regional and temporal expression of mitochondrial genes after CCI in female and male mice. Expression changes in mitochondrial-related genes following CCI in (A) female and (B) male mice across cortical regions (2, 4, 6 mm from epicenter) and hippocampus at 6, 12, 24, and 48 h post-injury. Circle color represents signed Z-score, which reflects the standardized difference between CCI and sham groups, accounting for both the magnitude of change and within-group variability (red, upregulation in CCI; blue, downregulation). Circle size indicates statistical significance; FDR values derived from Benjamini-Hochberg correction of Welch’s t-test p-values (larger circles = lower FDR). Black outlines mark FDR < 0.05. Genes grouped by functional category.

### Protein Markers of Mitochondrial Biogenesis and Quality Control After CCI

We next assessed changes in proteins representing functions like mitochondrial biogenesis, transcription, electron transport chain (ETC), and mitophagy. Protein abundance for PGC-1α, Tfam (A Mitochondrial transcription factor), Ndufs1, and p62 were quantified in cortex (2, 4, and 6 mm from epicenter) and hippocampus at 6h, 12h, 24h, and 48h post-CCI in both sexes (Figs.4-7; Supplementary Figure 1). Overall, CCI elicited region-restricted and sex-dependent alterations, with females showing earlier changes and males showing more delayed or blunted responses.

**FIG. 4.**
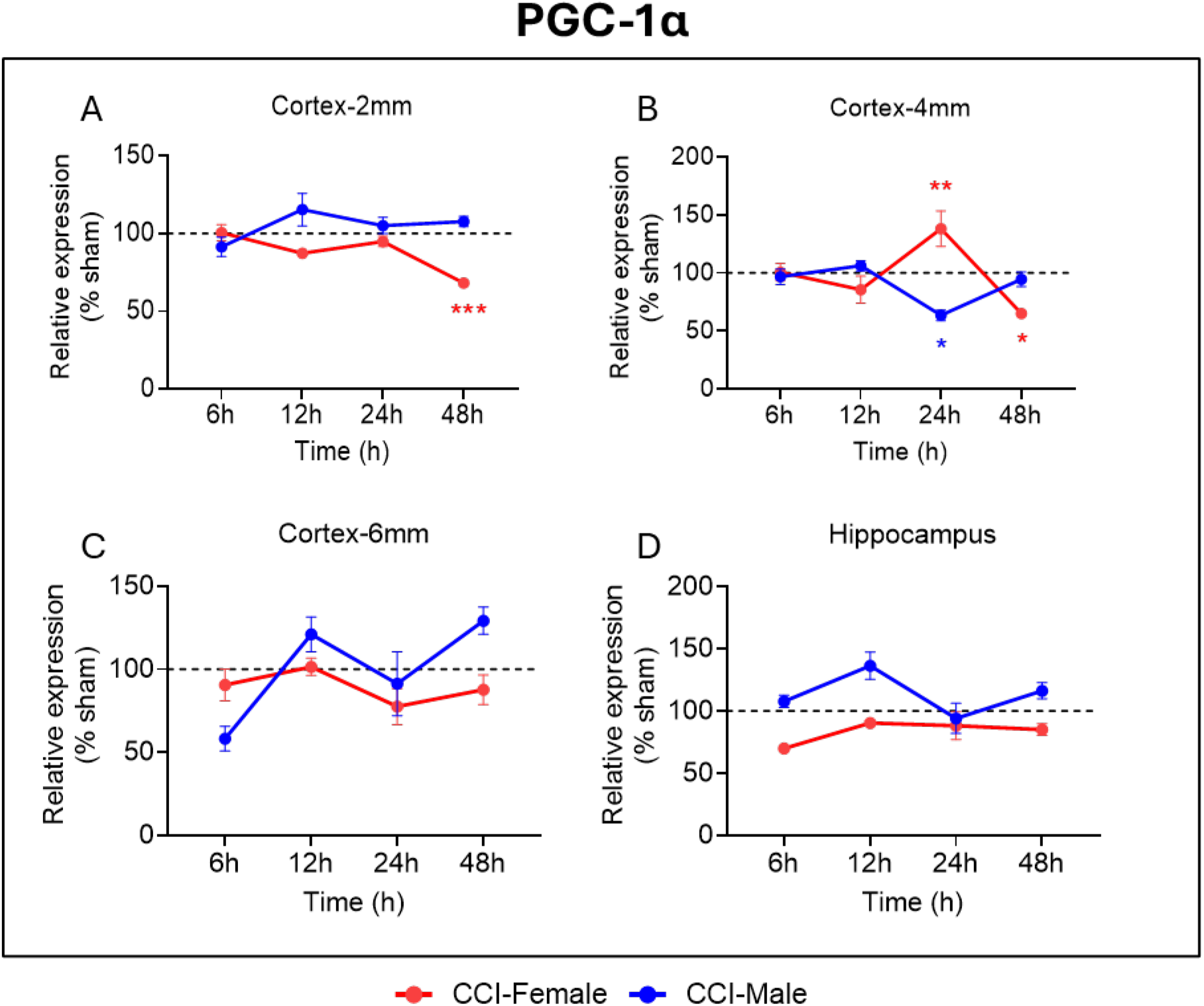
PGC-1α expression following controlled cortical impact (CCI) in female and male mice. PGC-1α protein levels were quantified in the **(A)** cortex-2 mm, **(B)** cortex-4 mm, **(C)** cortex-6 mm regions relative to the impact site, and **(D)** hippocampus at 6, 12, 24, and 48 h following CCI in female (red) and male (blue) mice. Data are expressed as relative expression (% of sham), with dotted horizontal lines indicating the sham reference level across all time points and regions. Data are presented as mean ± SEM, with individual biological replicates shown (n = 5–6 per group). Statistical significance is indicated as *p ≤ 0.05, **p ≤ 0.01, and ***p ≤ 0.001 compared with sham controls.

**FIG. 5.**
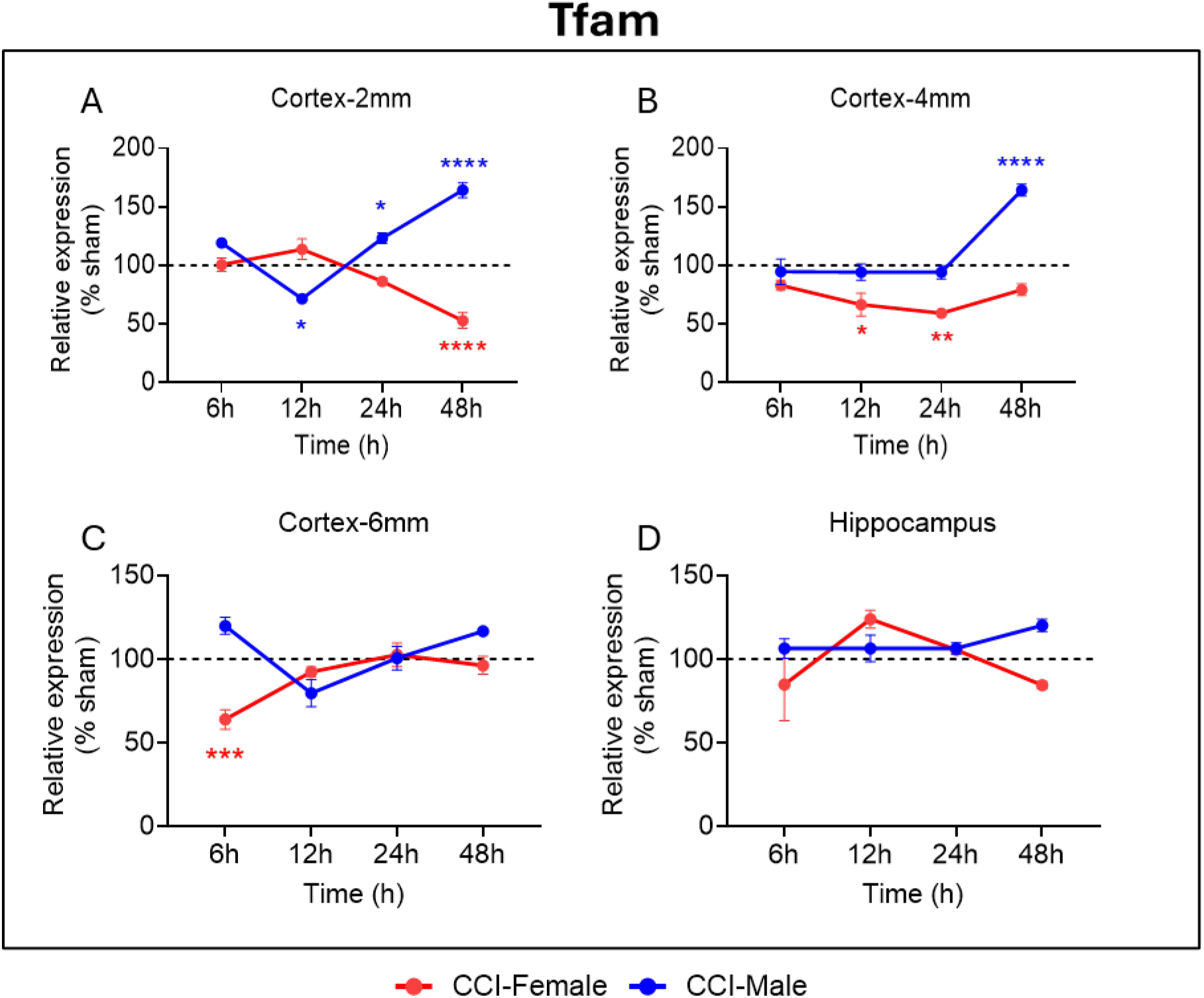
Tfam expression following controlled cortical impact (CCI) in female and male mice. Tfam protein levels were quantified in the **(A)** cortex-2 mm, **(B)** cortex-4 mm, **(C)** cortex-6 mm regions relative to the impact site, and **(D)** hippocampus at 6, 12, 24, and 48 h following CCI in female (red) and male (blue) mice. Data are expressed as relative expression (% of sham), with dotted horizontal lines indicating the sham reference level across all time points and regions. Data are presented as mean ± SEM, with individual biological replicates shown (n = 5–6 per group). Statistical significance is indicated as *p ≤ 0.05, **p ≤ 0.01, and ***p ≤ 0.001 compared with sham controls.

**FIG. 6.**
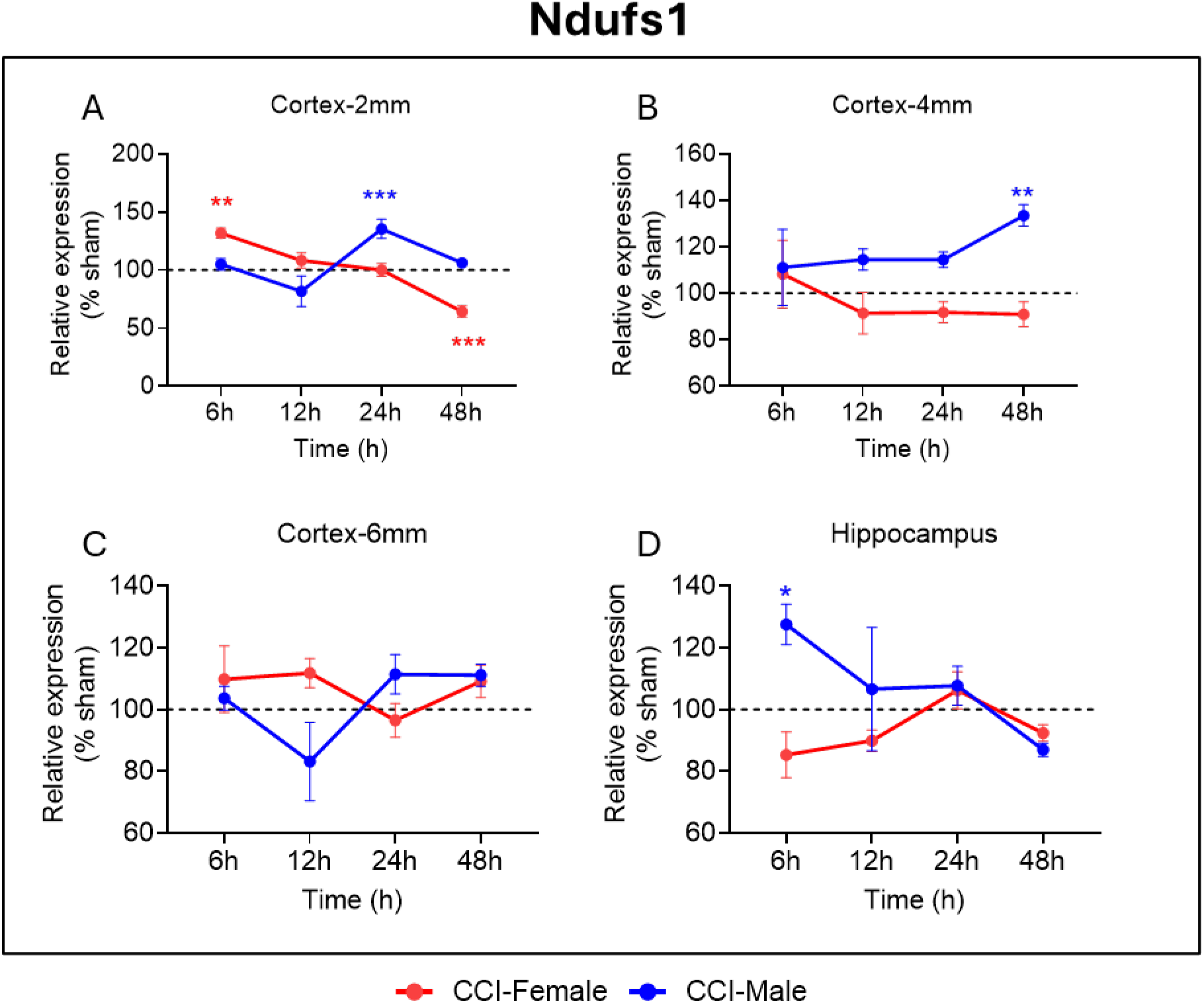
Ndufs1 expression following controlled cortical impact (CCI)) in female and male mice. Ndufs1 protein levels were quantified in the **(A)** cortex-2 mm, **(B)** cortex-4 mm, **(C)** cortex-6 mm regions relative to the impact site, and **(D)** hippocampus at 6, 12, 24, and 48 h following CCI in female (red) and male (blue) mice. Data are expressed as relative expression (% of sham), with dotted horizontal lines indicating the sham reference level across all time points and regions. Data are presented as mean ± SEM, with individual biological replicates shown (n = 5–6 per group). Statistical significance is indicated as *p ≤ 0.05, **p ≤ 0.01, and ***p ≤ 0.001 compared with sham controls.

**FIG. 7.**
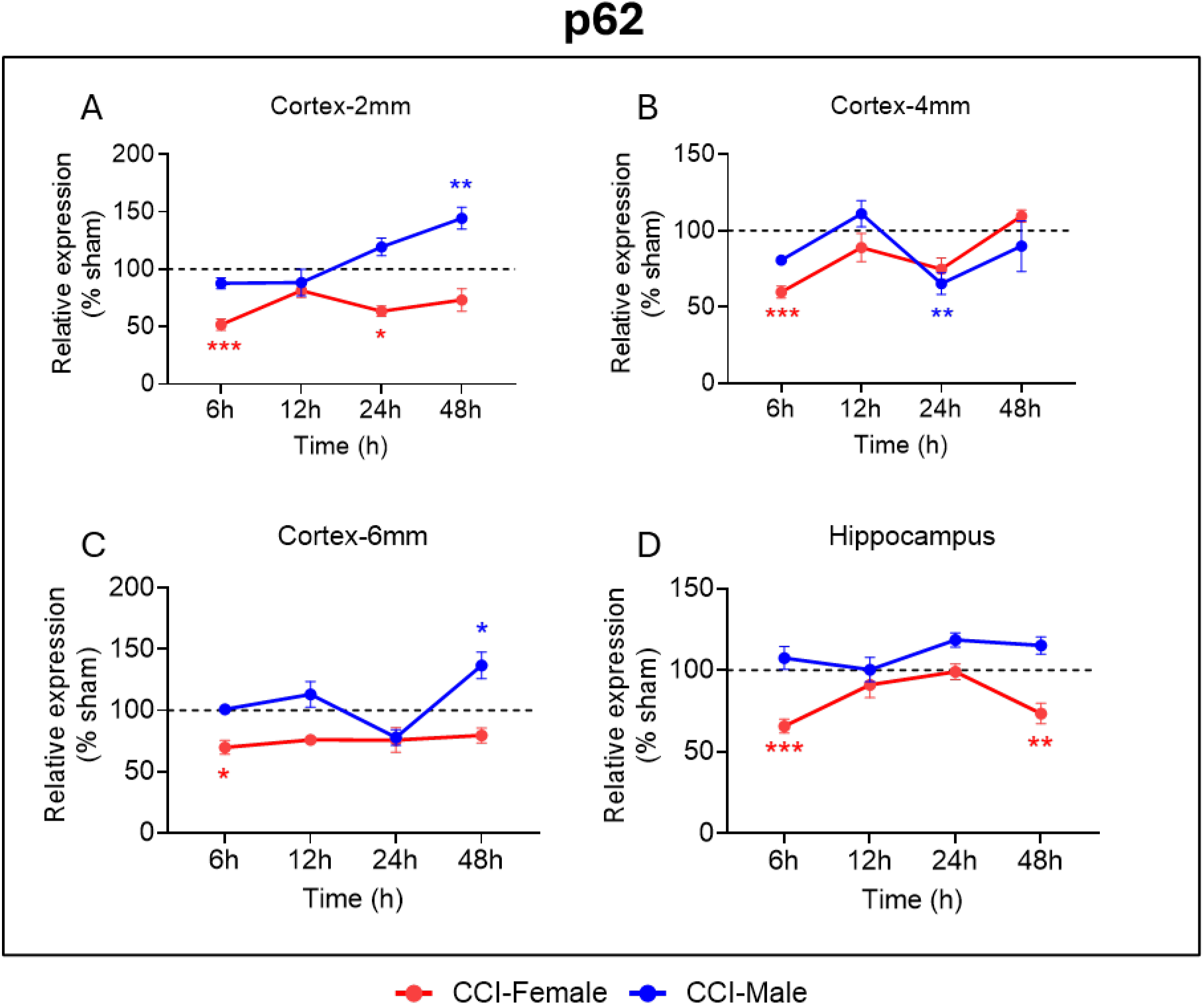
p62 expression following controlled cortical impact (CCI)) in female and male mice. p62 protein levels were quantified in the **(A)** cortex-2 mm, **(B)** cortex-4 mm, **(C)** cortex-6 mm regions relative to the impact site, and **(D)** hippocampus at 6, 12, 24, and 48 h following CCI in female (red) and male (blue) mice. Data are expressed as relative expression (% of sham), with dotted horizontal lines indicating the sham reference level across all time points and regions. Data are presented as mean ± SEM, with individual biological replicates shown (n = 5–6 per group). Statistical significance is indicated as *p ≤ 0.05, **p ≤ 0.01, and ***p ≤ 0.001 compared with sham controls.

### PGC-1α: Mitochondrial Biogenesis Master Regulator

PGC-1α protein levels showed region- and time-dependent alterations following CCI, with more pronounced effects in females (Fig.4). In female mice, PGC-1α was significantly reduced at 48 h in both cortex-2 mm (Fig.4A: F (3, 75) = 6.994; p = 0.001) and cortex-4 mm (Fig.4B: F (3, 75) = 1.258; p = 0.021), whereas earlier time points (6, 12, and 24 h) showed no significant changes. Interestingly, cortex-4 mm exhibited a brief increase in PGC-1α at 24 h before declining at 48 h (Fig.4B: F (3, 75) = 1.258; p = 0.006), suggesting a transient compensatory response. No significant alterations were observed in the cortex-6 mm (Fig.4C) and hippocampus (Fig.4D) region at any time point, indicating that, unlike other brain regions, PGC1a levels in the cortex-6 mm and hippocampus remain unaffected over time. In male mice, PGC-1α expression was largely stable across regions and time points, with a single significant decrease detected only at 24 h in cortex-4 mm (Fig.4B: F (3, 75) = 1.258; p = 0.015). No differences were observed in cortex-2 mm, cortex-6 mm, or hippocampus at any time. Overall, both sexes showed late-phase reductions in PGC-1α, particularly near the injury site (2 mm and 4 mm) and in the hippocampus, though females exhibited broader and more consistent suppression. These findings indicate sex-dependent vulnerability of mitochondrial biogenesis after CCI, with females showing a more pronounced and sustained decline.

### TFAM: Mitochondrial DNA Transcription and Maintenance Factor

TFAM protein expression demonstrated region- and sex-dependent alterations following CCI, with distinct temporal trajectories in male and female mice (Fig.5). In females, TFAM levels remained unchanged at early time points (6, 12, and 24 h) in the cortex-2 mm region, but a significant reduction emerged at 48 h (Fig.5A: F (3, 72) = 17.21; p = 0.0001), indicating a delayed impairment in mtDNA transcription and maintenance near the injury core. In the cortex-4 mm region, TFAM showed no change at 6 h but was significantly reduced at both 12 h (Fig.5B: F (3, 75) = 18.54; p = 0.021) and 24 h (Fig.5B: F (3, 75) = 18.54; p = 0.003), with expression returning to baseline by 48 h. In the cortex-6 mm region, TFAM exhibited a transient decrease at 6 h (Fig.5C: F (3, 78) = 4.281; p = 0.0008), which normalized at later time points. No significant alterations were detected in the hippocampus across the 48-hour period (Fig.5D). Collectively, the female profile suggests early transient TFAM suppression in peri-injury regions and a delayed decline at the injury core, reflecting impaired mtDNA regulatory capacity during both the acute and evolving stages of injury. Males displayed a more dynamic and compensatory pattern. TFAM levels showed a significant decrease at 12 h in the cortex-2 mm region (Fig.5A: F (3, 72) = 17.21; p = 0.013), mirroring one of the early female responses. Unlike females, however, males exhibited significant TFAM upregulation at both 24 h (Fig.5A: F (3, 72) = 17.21; p = 0.044) and 48 h (Fig.5A: F (3, 72) = 17.21; p < 0.0001) time points in cortex-2 mm. Similar increases were observed at 48 h in the cortex-4 mm region (Fig.5B; F (3, 75) = 18.54; p < 0.0001), along with a comparable upward trend in the cortex-6 mm region (Fig.5C), suggesting activation of late-phase mitochondrial biogenesis and mtDNA maintenance responses. As in females, hippocampal TFAM levels remained unchanged in male at all time points (Fig.5D). Altogether, TFAM expression patterns indicate opposing late-phase mitochondrial regulatory strategies between sexes, females show suppression, whereas males demonstrate delayed compensatory upregulation. These findings highlight fundamental sex-specific differences in mitochondrial genome regulation after TBI and suggest that males may initiate recovery-oriented biogenesis mechanisms that are diminished or absent in females.

### NDUFS1: Complex I Subunit of the Electron Transport Chain

NDUFS1 protein levels displayed clear sex- and region-dependent differences following CCI (Fig. 6). In females, NDUFS1 showed a transient early increase at 6 h in cortex-2 mm (Fig.6A: F (3, 75) = 1.345; p = 0.001), suggesting an acute compensatory upregulation of Complex I function in the peri-injury zone. At the injury-adjacent cortex-2 mm, NDUFS1 expression remained unchanged at 12 and 24 h but was significantly reduced at 48 h (Fig.6A: F (3, 75) = 1.345; p = 0.0004), paralleling the late suppression of mitochondrial ETC transcripts described earlier. At any time point, no significant alterations were observed in cortex-4 mm (Fig. 6B), cortex-6 mm (Fig.6C), and hippocampal NDUFS1 levels (Fig.6D) relative to sham, indicating preservation of mitochondrial Complex I in regions farther from the injury site. In males, NDUFS1 expression remained stable at 6 and 12 h in cortex-2 mm but showed a significant increase at 24 h (Fig.6A: F (3, 75) = 1.345; p = 0.0002) before returning to baseline by 48 h, reflecting a delayed but transient compensatory response. In cortex-4 mm, NDUFS1 levels were significantly elevated only at 48 h (Fig.6B: F (3, 77) = 7.735; p = 0.008), without changes at earlier time points. No significant differences were detected in cortex-6 mm (Fig.6C). Notably, the hippocampus exhibited a robust early increase at 6 h (Fig.6D: F (3, 76) = 2.249; p = 0.048), suggesting rapid activation of Complex I related mitochondrial support mechanisms in males. Collectively, these findings indicate that males exhibit delayed but compensatory upregulation of NDUFS1, particularly in peri-injury cortex and hippocampus, whereas females show an early transient increase followed by a pronounced late decline near the injury core. This pattern suggests sex-divergent mitochondrial stress responses, with females showing heightened vulnerability reflected by late Complex I suppression, and males demonstrating region-specific compensatory activation during the acute–subacute phase of injury.

### p62: Marker of Mitophagy and Autophagy Flux

p62 protein expression demonstrated clear sex- and region-dependent differences in mitophagy-related responses following CCI (Fig.7). Females showed a pronounced early decrease in p62 at 6 h across nearly all regions, including cortex-2 mm (Fig.7A: F (3, 75) = 21.87; p = 0.0002), cortex-4 mm (Fig.7B: F (3, 76) = 5.726; p = 0.0009), cortex-6 mm (Fig.7C: F (3, 78) = 11.76; p = 0.049),and hippocampus (Fig.7D: F (3, 76) = 17.85; p = 0.0002), indicating rapid activation of mitophagy and early clearance of damaged mitochondria following injury. p62 levels returned to baseline at 12 and 24 h in most regions, except for a transient reduction in cortex-2 mm at 24 h (Fig.7A: F (3,75) = 21.87; p = 0.0112) and a later decrease in the hippocampus at 48 h (Fig.7D: F (3, 76) = 17.85; p = 0.0085). This pattern suggests a short-lived but widespread autophagic response that resolves as acute mitochondrial stress diminishes. In contrast, males exhibited no early changes in p62 expression. Instead, significant increases emerged at 48 h in cortex-2 mm (Fig.7A: F (3, 75) = 21.87; p = 0.002) and cortex-6 mm (Fig.7C: F (3, 78) = 11.76; p = 0.011), consistent with delayed accumulation of autophagic cargo. This elevation may reflect impaired mitophagic flux or insufficient clearance of damaged mitochondria during the subacute phase of injury. Cortex-4 mm displayed a significant decrease at 24 h (Fig.7B: F (3, 76) = 5.726; p = 0.005), with no changes at 6, 12, or 48 h. No alterations in p62 were detected in the male hippocampus at any time point (Fig. 7D. Together, these findings indicate sex-divergent mitophagy dynamics after CCI. Females exhibit a rapid, early activation of mitochondrial clearance pathways, while males show delayed p62 accumulation, suggesting later-stage impairment or altered regulation of mitophagy. These opposing temporal patterns highlight fundamental sex differences in mitochondrial quality-control responses following TBI.

## Discussion

This study reveals that mitochondrial dysfunction after CCI is spatially graded, temporally dynamic, and profoundly sex-dependent, with females showing heightened variability at transcriptional and protein-regulatory levels despite relatively stable mtDNA content. Prior work established that mitochondrial dysfunction varies by injury severity, time, and region ^11,25-27^. Functional analyses demonstrated time-dependent changes in state III respiration and respiratory control ratios after CCI ^28^, and using Dendra2-labeled mitochondria, our lab showed increased mitochondrial fragmentation in the injured cortex ^29^. Active hippocampal neurodegeneration, assessed by silver staining, peaks at 48 hours post-CCI and largely abates by 7 days ^30^. Collectively, these studies indicate that acute and subacute mitochondrial dysfunction reflects a dynamically changing bioenergetic environment. However, prior work examined these variables largely in isolation. By integrating mtDNA-CN, gene expression, and protein markers across multiple cortical regions, hippocampus, and both sexes within a single study, we define how spatial, temporal, and sex-dependent factors interact during the first 48 hours after injury, a critical window for mitochondria-targeted therapeutics ^5^.

MtDNA plays a critical role in driving pathophysiology of TBI. MtDNA-CN varies widely across tissues depending on developmental stage and pathophysiological conditions ^31^. mtDNA-CN and heteroplasmy are key determinants of degenerative and neurodegenerative conditions, including TBI ^32,33^. Cerebral mitochondrial dysfunction influences the peripheral blood mtDNA content which could be used as a biomarker in TBI ^34^. Increasing evidence suggests mtDNA-CN as an important marker of mitochondrial mass and therefore cellular energy homeostasis. Our results demonstrate that acute changes in mtDNA-CN are centered around the injury site within a 2mm diameter at 6h and the trend was maintained till 24h (Fig.2A). Although the overall trend remained decreased, spatial and temporal gradients became less pronounced with increasing distance from the injury site and at later post-injury intervals (Fig.2B-C), as well as across all time points in the hippocampus (Fig.2D). When accounting for sex differences, the overall trend in mtDNA-CN remained consistent, however, male mice displayed a more pronounced decline than female mice. Overall, the changes correlate with the acuteness and proximity to the site of the injury which contrast with the transcriptional and translational level mitochondrial regulations. This indicates that initial changes in mitochondria correlate with the early primary injury and structural loss of mitochondria which is compensated for overtime that may or may not be able to restore the bioenergetic homeostasis. Substantial research on transcriptional and translational regulation of mitochondrial function has been carried out but very few studies have examined the spatial and temporal dynamics of mtDNA-CN or the mechanisms underlying these regulatory changes following TBI ^35^. The observed acute decline in mtDNA-CN is likely driven by mtDNA damage, increased mitophagy, and impaired mitochondrial biogenesis ^33,36^. In addition to localized cellular mitochondrial alterations, other factors likely contribute to mtDNA variability, including acute necrosis and rapid immune cell infiltration at the injury site. In contrast, longer-term increases in mtDNA-CN have been reported after TBI which may reflect compensatory mitochondrial adaptations aimed at restoring bioenergetic balance ^35^. Although the mtDNA-CN is widely accepted as a marker for mitochondrial mass, no study demonstrates the relation between the mitochondrial state or health with its relative DNA content. Given these findings, further research is necessary to elucidate the cell-specific compensatory mechanisms of mtDNA-CN and the subsequent regulation of mitochondrial function following brain injury..

A panel of 20 genes related to mitochondrial metabolism was selected for expression analysis (Table 1). Based on functional roles, these genes were categorized into mitochondrial dynamics, mitochondrial function, and transcriptional regulation groups. Gene expression profiling revealed pronounced sex-, time-, and region-dependent vulnerability. Notably, female mice accounted for most transcriptional changes, exhibiting early upregulation of mitochondrial function and dynamics-related genes in the injury-proximal cortex (2-4 mm) within the first 12 hours post-injury, followed by marked downregulation at 48 hours across multiple pathways, including oxidative phosphorylation, mitochondrial fusion/fission, biogenesis, and transcriptional regulation Although the overall trend remained decreased, spatial and temporal gradients became less pronounced with increasing distance from the injury site and at later post-injury intervals (Fig.2B-C), as well as across all time points in the hippocampus (Fig.3A). This biphasic pattern suggests an early compensatory response that becomes unsustainable as injury-related stress accumulates. Sex-specific outcomes in TBI remain heterogenous mainly due to differences in the injury severity and measured outcomes; however, most preclinical studies report improved early outcomes in females ^37^, likely due to the neuroprotective effects of estrogen and progesterone. A striking early increase in mitochondrial dynamics genes, along with structural and regulatory genes in female mice, indicates a mitochondrial remodeling response in the injury core (2 mm) and immediate perilesional region (4 mm) during the 6–12 h window. Females exhibited significant upregulation of Prkaa2, accompanied by increased expression of Pink1, Park2, Mfn1, Fis1, and Dnm1, consistent with adaptive mitochondrial quality control and remodeling. Concurrent upregulation of mtDNA-encoded structural genes (Nd1, Cox1, and Atp6) suggests enhanced compensatory bioenergetic activation, which may explain the relatively smaller decline in mtDNA-CN observed in females. Despite being anatomically distant from the impact site, the hippocampus displayed the most pronounced late gene suppression in females at 48 hours, highlighting the widespread effects of secondary injury cascades (Fig.3A). This delayed vulnerability could be one of the reasons for heterogeneity across injury response and the lack of sustained sex-specific differences in long-term tissue sparing ^38^. In contrast, males exhibited minimal early transcriptional perturbation, suggesting reduced mitochondrial plasticity at transcriptional levels during the acute post-injury phase (Fig.3B). The hippocampal sex differences align with our prior findings showing delayed bioenergetic impairment in non-synaptic mitochondria in females but not males ^4^. The absence of similar differences in synaptic mitochondria suggests that these sex-specific effects may be driven by differential activation and infiltration of glial populations. Together, these findings underscore the importance of investigating mitochondrial responses to TBI in a cell type– and region-specific manner.

Next, we examined key protein-level changes in the same tissue homogenates to further evaluate sex-divergent mitochondrial responses (Fig.4-7; Supplementary Fig.1). We focused on four proteins: PGC-1α (Fig.4), TFAM (Fig.5), NDUFS1 (Fig.6), and p62 (Fig.7). PGC-1α is a central regulator of mitochondrial homeostasis, controlled by multiple signaling pathways and widely implicated in neurodegeneration and brain injury ^10,39^. No significant changes in PGC-1α were observed up to 12 hours post-injury. In the 2 mm cortex, PGC-1α decreased at 48 hours in females but remained unchanged in males (Fig.4A). In the 4 mm cortex, males and females exhibited divergent responses at 24 hours, with decreased PGC-1α in males but increased levels in females (Fig.4B), likely reflecting the earlier transcriptional upregulation observed at 12 hours in females but not males. By 48 hours, PGC-1α declined in females but remained unchanged in males (Fig.4B), indicating an overall reduction in PGC-1α signaling in the injured female cortex. TFAM is a multifunctional mitochondrial protein involved in mtDNA transcription, replication, and maintenance ^40^, and is indirectly regulated by PGC-1α via NRF1/2 and ERRα. In females, TFAM levels decreased in the 2 mm cortex at 48 hours (Fig.5A) and in the 4 mm cortex between 12 and 24 hours (Fig.5B). In contrast, TFAM levels increased in males at 48 hours in both 2mm (Fig.5A) and 4mm (Fig.5B) regions. This pattern suggests that in females PGC-1α upregulation does not result in increased downstream TFAM, instead TFAM expression remained low at 48h time points. In males, upregulation of TFAM at 48h despite stable PGC-1α levels suggest a possible alternative regulation, such as altered PGC-1α nuclear localization or delayed compensatory signaling. The increase in TFAM in males may also represent a compensatory response to greater mitochondrial loss, consistent with the more pronounced decline in mtDNA-CN. NDUFS1, a core subunit of complex I of the electron transport chain ^41^, exhibited trends like TFAM. In females, NDUFS1 decreased at 48 hours in the 2 mm region, whereas males showed increased expression at 24 hours in the 2 mm cortex (Fig.6A) and at 48 hours in the 4 mm cortex (Fig.6B). These findings suggest a more delayed but potentially compensatory mitochondrial response in males relative to females. p62, an established marker of autophagy pathway engagement ^42^, showed early reductions in females at 6 hours across all 4 regions, with levels remaining suppressed through 48 hours. In contrast, males exhibited increased p62 expression at 48 hours in the 2 mm (Fig.7A) and 6 mm (Fig.7B) cortex. Changes in p62 may reflect either enhanced autophagic activity or impaired downstream autophagic flux, which will require further investigation. Nevertheless, the sex-specific p62 trajectories align with broader protein-level differences observed across mitochondrial pathways.

Key limitations of this study include its restriction to the first 48 hours post-injury, precluding conclusions about long-term mitochondrial adaptation or chronic neurodegeneration. The analyses were performed in bulk tissue homogenates, limiting cell type–specific resolution and making it difficult to distinguish neuronal from glial mitochondrial responses. Additionally, mechanistic links between sex hormones and the observed transcriptional versus protein-level regulation were inferred but not directly tested. Future studies should determine how cell type-specific mitochondrial alterations in neurons, astrocytes, microglia, and infiltrating immune cells shape regional metabolic outcomes after injury. Longitudinal investigations extending into subacute and chronic phases are needed to establish whether early compensatory responses predict long-term recovery or degeneration. Direct assessment of mitochondrial health, including respiration, ROS production, mitophagic flux, and mtDNA integrity will clarify the functional significance of mtDNA-CN changes. It will also be critical to define how estrogen and progesterone signaling mechanistically regulate mitochondrial biogenesis and quality control pathways after TBI. Finally, evaluating how these sex-dependent mitochondrial trajectories influence therapeutic responsiveness will inform precision-based, mitochondria-targeted interventions

This study helps define phase-specific therapeutic windows for mitochondria-targeted interventions after TBI. The early 6–12-hour period, marked by acute mtDNA decline and active mitochondrial remodeling, represents a window for therapies that stabilize mitochondria, reduce ROS, or enhance mitophagy before irreversible damage occurs. By 48 hours, transcriptional suppression (especially in females) and delayed compensatory protein responses (notably in males) suggest a shift toward secondary bioenergetic failure, where strategies promoting mitochondrial biogenesis or metabolic support may be more effective. The spatial gradient of injury further supports targeted regional delivery, while sex-dependent differences indicate that optimal timing and mechanism of intervention may need to be sex-specific. Overall, the current findings and the future studies in this direction will enable rational alignment of drug mechanism with injury stage for precision mitochondrial therapy design.

In summary, our findings reveal that mitochondrial responses to traumatic brain injury evolve along distinct spatial, temporal, and sex-dependent trajectories during the early post-injury period. While mtDNA content shows only localized changes near the injury core, transcriptional and protein-level regulation of mitochondrial pathways demonstrates pronounced and divergent responses between males and females. Females exhibit early mitochondrial remodeling followed by widespread suppression of mitochondrial programs, whereas males display delayed but potentially compensatory regulatory responses. These results highlight mitochondrial regulation as a dynamic and context-dependent component of secondary injury and underscore the importance of incorporating spatial resolution and sex as key biological variables when designing mitochondria-targeted therapeutic strategies for TBI.

### Transparency, Rigor, and Reproducibility Statement

All data are reported as mean ± SEM with individual biological replicates shown where applicable. Reagents, antibodies, and experimental conditions are described in detail to enable replication. Raw data supporting the findings of this study are available from the corresponding author upon reasonable request.

## Acknowledgements

We are thankful to Frances I. Meredith, Laboratory Technician in Patrick Sullivan laboratory at University of Kentucky, for maintaining animal inventory and ordering supplies required for the study.

## Author Contributions

P.G.S., H.J.V., and W.B.H. conceptualized the study. Resources were provided by P.G.S., who also contributed to the review and editing of the manuscript. C.D.P. and H.J.V. were responsible for original draft preparation, data analysis, and figure design. H.J.V., P.P., E.Z.M., and G.V.V. performed the CCI surgeries and associated assays. A.D.B. contributed to the transcriptomic data analysis, visualization, and manuscript editing. All authors have read and agreed to the published version of the manuscript.

## Supplementary Material

Supplementary Figure S1

