## Supplemental Figure for "Spatial, temporal and sex specific mitochondrial dynamic changes in severe controlled cortical impact mouse model of traumatic brain injury": Vekaria et al_Supplyment_BioRxiv.pdf

**Supplementary Material**

Supplementary Figure S1

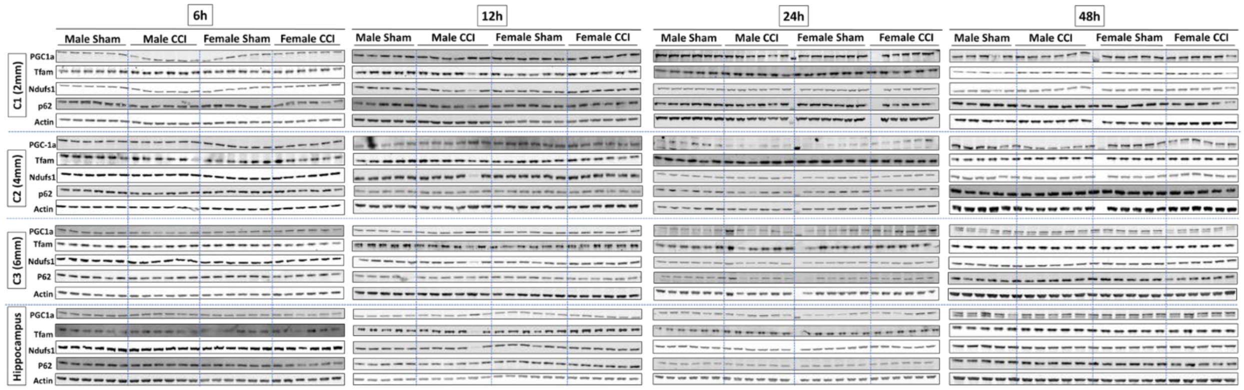


**Figure Legends:**

**Supplementary FIG.1. Immunoblot validation of mitochondrial protein expression across cortical regions and hippocampus following controlled cortical impact (CCI) in female and male mice.** Supplementary Figure 1 shows the raw immunoblot images corresponding to the quantitative analyses presented in Figures 4–7. Protein levels of PGC-1α (master regulator of mitochondrial biogenesis), Tfam (mitochondrial transcription factor A), Ndufs1 (Complex I subunit), p62/SQSTM1 (mitophagy adaptor protein), and β-actin (loading control) were assessed in a spatially resolved (cortex 2 mm [C1], 4 mm [C2], 6 mm [C3] from the injury epicenter, and hippocampus) and temporally resolved (6, 12, 24, and 48 h post-CCI) manner in both female and male mice. For each protein, representative blots are shown from sham (n=5-6) and CCI (n=5-6) conditions across all regions and time points.
